# Scalable clustering with supervised linkage methods

**DOI:** 10.1101/2021.08.01.454697

**Authors:** James Anibal, Alexandre G. Day, Erol Bahadiroglu, Liam O’Neill, Long Phan, Alec Peltekian, Amir Erez, Mariana Kaplan, Grégoire Altan-Bonnet, Pankaj Mehta

## Abstract

Data clustering plays a significant role in biomedical sciences, particularly in single-cell data analysis. Researchers use clustering algorithms to group individual cells into populations that can be evaluated across different levels of disease progression, drug response, and other clinical statuses. In many cases, multiple sets of clusters must be generated to assess varying levels of cluster specificity. For example, there are many subtypes of leukocytes (e.g. T cells), whose individual preponderance and phenotype must be assessed for statistical/functional significance. In this report, we introduce a novel hierarchical density clustering algorithm (HAL-x) that uses supervised linkage methods to build a cluster hierarchy on raw single-cell data. With this new approach, HAL-x can quickly predict multiple sets of labels for immense datasets, achieving a considerable improvement in computational efficiency on large datasets compared to existing methods. We also show that cell clusters generated by HAL-x yield near-perfect F1-scores when classifying different clinical statuses based on single-cell profiles. Our hierarchical density clustering algorithm achieves high accuracy in single cell classification in a scalable, tunable and rapid manner. We make HAL-x publicly available at: https://pypi.org/project/hal-x/

## Introduction

Technological advances have made it possible to collect huge single–cell datasets with numerous features *e.g.* transcriptomic profiles (scRNAseq) and/or protein expression (Flow cytometry, CyTOF, CyTEK or CiteSEQ). The ability to cluster such large, high-dimensional datasets is important for a variety of data-intensive fields ranging from biology to data mining. For this reason, there is a crucial need for fast, tunable and scalable clustering algorithms that work well in a high-dimensional setting on limited computational resources and that are at the same time reliable and robust. Clustering large high-dimensional datasets presents several challenges. Clustering in high dimensions suffers from the “curse of dimensionality” [1, 2]. The number of parameters in model-based clustering methods explodes and it becomes difficult to obtain accurate density estimates, a crucial ingredient in density-based clustering algorithms. Since many features are irrelevant or noisy, it is hard to construct meaningful similarity measures [3]. In order to circumvent these problems, one often needs to perform dimensional reductions and/or construct large similarity graphs, but both of these methods are computationally prohibitive for large datasets. Another difficult component of clustering is using the data to automatically learn the number of ideal clusters. Finally, arguably the most controversial and subjective part of clustering is validating clustering assignments when no ground-truth labels are available.

In scientific research, clustering algorithms provide many advantages and are widely used. These algorithms are easy to use, readily explainable, and do not require any form of annotated data. Nonetheless, clustering does have a significant downside: the requirement of prior knowledge about the data. A common task in modern biology is to cluster individual cells into distinct cell populations based on measurements in a high-dimensional feature space (mRNA or protein levels). Here, we focus largely on mass cytometry data (CyTOF) which allow measurements of more than 30 features, often for tens of millions of cells in a single sample [4, 5]. When performing exploratory analysis of unsupervised data, researchers often do not know the specificity (*i.e.* depth of clustering) needed to derive biologically-meaningful populations. A rare yet important population of cells may be lost when there are insufficient clusters. Conversely, it may be beneficial to study larger, well-understood populations that yield immediately actionable insights with robust statistical features.

To perform this type of comprehensive analysis, multiple sets of clusters must be generated across a broad spectrum of specificity. For currently available methods (i.e., Parc, Phenograph), this requires training multiple models with different parameters to control the number of clusters. In this scenario, time and/or memory complexity of these algorithms generate a significant problem, particularly in the area of cluster prediction. In the case of training, the solution is to perform dimensional reduction of the feature space (e.g. PCA the features) as well as possibly downsampling the data [5, 6].

In the context of single-cell analysis, there are disadvantages to downsampling the dataset. We can identify generalizable cell populations with downsampled data, but we cannot identify nuanced perturbations within populations. Statistically significant features may be lost in the downsampling. For example, we can attempt to identify biomarkers by training a supervised model for separating healthy cells from diseased cells using only specific clusters. Intuitively, training on a small downsample will lead to severe overfitting, as the subtle differences are likely to be statistical noise and not be representative of the entire dataset. However, to create large, population-specific datasets, we must parse the full dataset based on phenotype, which requires annotations. Here, the apparent solution is to train a clustering model on a downsampled dataset and predict the labels of any remaining data or new data that is generated from additional samples. However, many of the current algorithms are transductive methods: the underlying model changes when considering a new data point. As such, Parc and Phenograph lack the ability to predict labels for newly acquired data points. HDBSCAN [7], allows for label prediction after the model has been trained by estimating the position of the new point within a fixed hierarchical tree. To do this, the feature space of the test data must be reduced to match the training data. However, dimensionality reduction on a 25.10^6^×40 matrix (a typical dataset acquired by mass cytometry [5]) will overwhelm the computing resources available to most biologists. Furthermore, prediction becomes more computationally expensive as the amount of training data is increased. This is because labeling is performed by nearest neighbour search rather than inputting the data into a learned function.

In this paper, we introduce a hierarchical density clustering algorithm (HAL-x) that uses supervised linkage methods to overcome these challenges. Our algorithm builds upon the idea that clustering can be viewed as a supervised learning problem where the goal is to predict the “true class labels” from data [8–10]. Unlike these earlier works that rely on the ideas of cluster stability, we operationalize this concept by training an expressive supervised learning model (*i.e.*, support vector machines, random forests) to evaluate potential cluster assignments and by using out-of-sample performance as a metric of clustering goodness. By training a sufficiently expressive supervised model, we can ensure that the predictive power is limited by the reproducibility of our clustering assignments and not by the choice of classifier [11]. Finally, we emphasize that using *out–of–sample* performance for clustering can be thought of as a generalization of density clustering to high-dimensional space, where direct density estimation is extremely hard. We empirically demonstrate that out-of-sample performance of a classifier trained on cluster labels captures intuitive ideas of good clustering. Additionally, such a robust clustering generalizes well and enables the analysis of large biological datasets.

The output of our algorithm is a predictive classifier that encodes our clustering assignments. Our work makes 3 primary contributions to unsupervised learning:

1. Since we can easily predict labels with supervised classifiers, we can extensively down-sample our data when performing computationally expensive tasks such as dimensionality reduction and the construction of similarity matrices (tasks which scale as O(n^2^) or O(n log n)).
2. Through the use of inductive learning methods, we can predict new data points at a minimum computational cost by simply applying the trained classifier on the full dataset. Moreover, we train these classifiers directly on unreduced data (i.e. without a dimensional reduction step), removing the need for expensive reduction of the test data.
3. Our model stores the classifiers corresponding to different out-of-sample error values. Thus, we can obtain multiple sets of cluster labels with a single model. We have converted our model into a simple python package such that users can easily generate multiple clustering, simply by specifying the desired value for the minimum out-of-sample error (intuitively, more subtle clusters lead to high error values).

HAL-x removes the need for pre-processing the entire dataset or training a model on the entire dataset. Moreover, HAL-x allows rapid, scalable prediction of new points. Finally, a single trained HAL-x model can generate multiple clusterings at varied depths to account for the specificity/sensitivity trade-off. Our results show that supervised linkage methods have great potential in analyzing high-dimensional data with limited computational resources that are typically available to experimental biologists (i.e., 8 core laptop computers).

## Related Work

Clustering has been extensively studied in the statistics and machine learning literature (see [3] for full review). Prominent methods for clustering include density-based methods [12, 13], spectral clustering methods [14], and model-based clustering [2]. In the high-dimensional context, several algorithms have been developed that seek to rely on clustering on a subspace of features [15]. Other methods rely on dimensional reduction methods such as Principal Component Analysis (PCA) and Local Discrimination Analysis (LDA) in conjunction with K-mean clustering (often in an iterative manner) [16–18]. There has also been considerable performance on extending methods that generalize manifold learning and spectral clustering to a high-dimensional setting [19, 20]. Several algorithms have been proposed for clustering in high-dimensions by first projecting the data using non-linear embeddings using such as t-SNE and then performing clustering in the low-dimensional space [21]. A drawback of these approaches is that low-dimensional representations can compress the data in such a way that spurious clusters may form in the low-dimensional representation [22], making it extremely challenging to identify true clusters from artifacts of visualizations. Additionally, there exists specialized methods developed specifically for high-dimensional biological data such as Phenograph [5], others, etc. However, these generally tend to require extensive computational resources to scale to large datasets.

More recently, state-of-the-art results were obtained by the DEC algorithm which combines deep embeddings with k-means clustering on the compressed, encoded representation [23]. The encoder weights and the means of the k-clusters were fine–tuned using a clever stochastic gradient descent–based training procedure. A particularly attractive aspect of this algorithm is that the cluster labels for new points can be calculated quickly by using the trained encoder to map points to the embedding space and finding the closest cluster center. However, an important drawback of this approach, shared by all k-means based approaches, is that it requires choosing the number of clusters by hand. This is often difficult to determine ahead of time for complex, high-dimensional data and retraining the deep encoder for different k can be time-consuming. Here we show how one can combine powerful non-linear embedding techniques, density clustering, and supervised methods that enforce self-consistency of cluster labeling to build a fast, accurate high-dimensional clustering algorithm that scales well to large datasets.

## Out-of-sample error captures clustering performance

Consider the problem of clustering n points 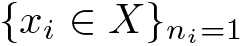 that live in a d-dimensional feature space. A (hard) clustering *C*(*x*) maps each point in the feature space to a set of discrete labels in a set *K*, with the cardinality of *K* equal to the number of possible cluster labels |*K*| = *k*. For example for k-means clustering, given a set of k-centroids 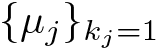, the cluster labels are determined by labelling each data point by the closest cluster centroid. Empirical and theoretical arguments suggest that meaningful clusterings *C*(*x*) should not differ much when learned using different subsets of the data and different measure of stability have been proposed to quantify this idea [8–10, 24]. Here, we extend this idea to argue that good clustering *C*(*x*) are maps that should generalize well. That is, given a clustering assignment generated on a subset of the data (a training data set), one should be able to accurately train a classifier to predict the cluster labels on unseen data (a test set). An important caveat is that the classifier we train should be expressive enough so that the predictive power of a classifier is limited by the properties of the clustering map rather than the classifier. It was recently shown that large, neural networks can achieve zero training error on random labels [11], suggesting that it is always possible to find such a classifier even for complex data. In practice, we have found that any reasonably powerful supervised learning algorithm (kernel-based SVMs, Random Forests, small neural networks) are sufficiently powerful to assess the goodness of clustering.

The intuition for using generalizability as a criteria is straightforward: clustering labels should reflect the underlying structure of the probability distribution from which the data is generated and we should be able to train an expressive classifier to learn this structure. Despite its intuitive appeal, it is extremely difficult to offer a rigorous theoretical justification for this idea. For this reason, we empirically tested whether having a low generalization error correlated with intuitive ideas of a good clustering. We started by considering simple low-dimensional datasets designed to highlight the caveats and strengths of standard clustering methods [see scikit-learn]. For each dataset, we performed clustering using different clustering methods (DBSCAN, Spectral clustering, k-means, Meanshift) with varying parameters (see supplementary materials for the parameters used). The results are presented in Figure 1, where we plotted the out-of-sample accuracy of the trained classifier against the comparison to the ground-truth labels. The latter is quantified by using the weighted F1-score which uses the Hungarian algorithm for the optimal cluster matching and equally weights small or large clusters. As can be seen in the figure, clusterings with high F1-scores also tend to have out good of sample accuracy confirming that out-of-sample accuracy is a reasonable proxy for good clustering.

**Fig 1.**
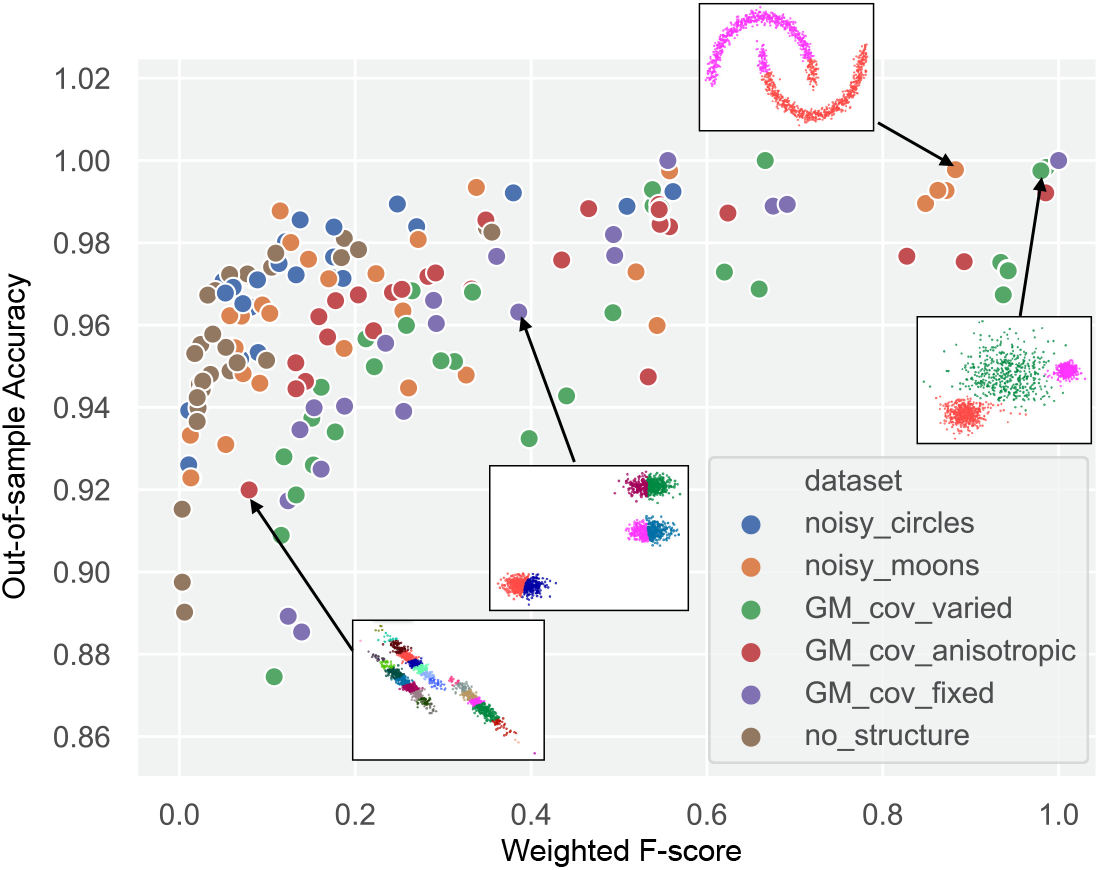
Out-of-sample accuracy vs. weighted *F*-score (comparison to ground-truth) for various clustering algorithms. We used out-of-the-box clustering algorithms *DBSCAN*, *spectral clustering*, *K*-means and *Meanshift* with various hyper-parameters and clustered various two-dimensional datasets taken from scikit-learn (see supplementary materials for more details). The out-of-sample accuracy is computed by training a support vector machine classifier with radial basis function kernels on the clustering labels provided by the clustering algorithms. We use a 80/20 train/test split to train the classifiers and evaluate the out-of-sample error.

## Methods

HAL-x has four major components that allow fast clustering of very large, high-dimensional datasets (Figure 2). First, HAL-x applies the t-SNE algorithm to reduce the dimensions on a down-sampled portion of the data. Second, HAL-x uses an approximate nearest neighbors algorithm and kernel density estimation to identify “pure clusters” in regions above a specified density threshold. Third, HAL-x defines an extended density neighborhood for each pure cluster, identifying spurious clusters that are representative of the same density maxima and should be merged. Fourth, HAL-x builds a sparsely connected K-nearest neighbor (KNN) graph that connects each cluster with the *k* most similar clusters. The edges of the graph are labeled with the accuracy of a supervised classifier (i.e. SVM, random forest) that is trained to separate unreduced data points taken from the two clusters in the original high-dimensional feature space. The clusters with the lowest edge-score (lowest classification accuracy) are then merged. An overview of the basic workflow is shown in Fig. 2. Runtime analysis and a pseudocode representation of the HAL-x algorithm can be found in the supplementary materials.

**Fig 2.**
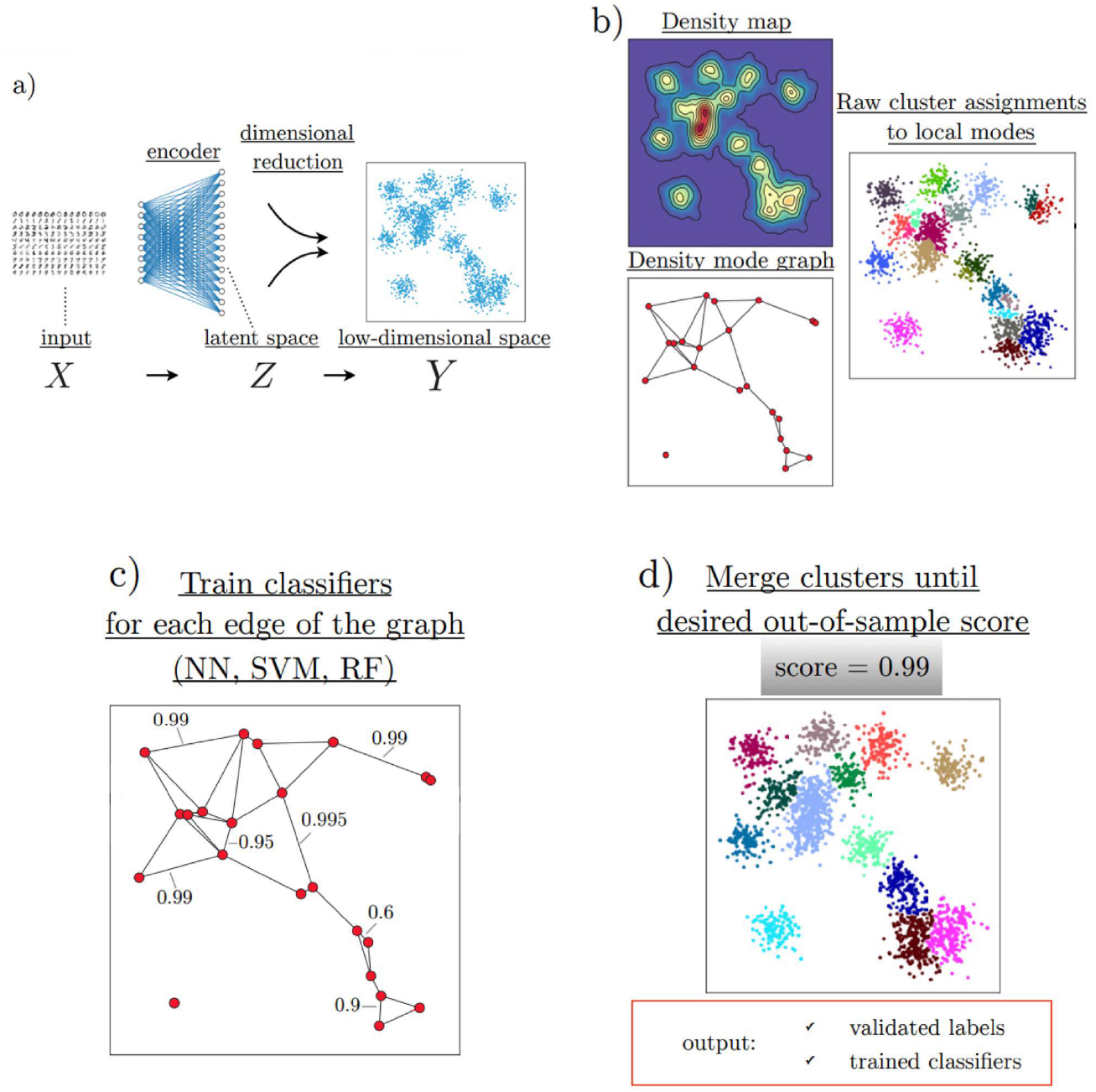
Overview of the hierarchical agglomerative learning approach. (a) For high-dimensional inputs, dimensional reduction (preferably preceded by an encoding to a latent space with high-level features, see [23] for instance) is necessary to obtain reliable density estimates. (b) In low-dimensional spaces, density maps can be easily computed. Initial clusters are selected to be the neighborhood of the density modes (see [13]). (c) A *k* nearest-neighbor graph is constructed by measuring similarity via the *out-of-sample accuracy* by training classifiers in the original high-dimensional feature space: each node represents an individual cluster and each edge has an associated weight given by the accuracy of the classifier. (d) Nodes are successively merged by pairs following the procedure until a desired out-of-sample accuracy is reached. The end result consists of an interpretable hierarchical classifier and robust clustering assignments. The classifier can be used to predict the labels of new data and potentially identify outliers.

### Dimensionality Reduction

HAL-x is designed to cluster datasets with up to 100 million points embedded in a 50+ dimensional space (typical datasets collected in mass cytometry measurements). To do this, the data must first be projected into a low-dimensional plane. HAL-x uses the z-score technique to normalize a down-sample of the data (the size of the down-sample is adjusted depending on the dimensionality of the dataset). This down-sample is used for the application of Fast Fourier Transform-accelerated Interpolation-based t-stochastic neighbor embedding (fitSNE) [25]. The application of fitSNE maps the data into a two-dimensional plane. t-SNE is especially well-suited for density clustering methods since it preserves local ordination in the data, while repelling points that are far away.

A drawback of t-SNE is that one must settle on a particular metric (usually Euclidean) to compute the initial similarity matrix before learning the embedding. This is problematic for data with highly correlated features and that have underlying labels invariant under global transformations (e.g. for images: translations and rotations). In order to alleviate this problem and generalize our approach to a broad class of high-dimensional datasets, we can also preprocess the input using an encoder if necessary. As in [27], we train our encoder by training a deep autoencoder and then discarding the decoder. The underlying intuition of the approach is to have the encoder learn high-level invariant features of the data in order to increase (conversely decrease) the similarity between data points belonging to similar clusters, improving the ability of HAL-x to capture nuance.

### Identification of Pure Clusters

HAL-x identifies an initial set of “pure”, high-density clusters that form the starting point for agglomerative clustering. This is done by performing density clustering in the low-dimensional embedding space using a variation of the algorithm proposed in [13]. In particular, cluster centers are identified by finding isolated high-density points, with nearby neighbor mapped to the closest cluster center.

To estimate local densities, HAL-x performs an approximate nearest neighbors search via the locality sensitive hashing (LSH) search algorithm with random projections. For each point, the number of neighbors is scaled linearly with the number of points in the down-sampled dataset.

Once HAL-x has identified the neighborhood of each point, a Gaussian kernel density estimate is used to determine if a point has the highest density in the neighborhood. Eq. 1 shows the relation that holds true for the probability of a point existing at *x* given a finite number of samples.

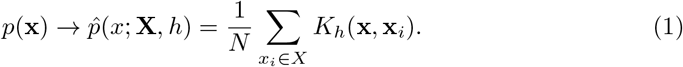

However, the bandwidth parameter, which can be thought of as a regularization parameter and sets how wide each kernel should be, must be determined. We do this using a cross-validated maximum likelihood estimation. We start by splitting the samples into a training and a testing set *X* = (*X*_train_, *X*_test_). Using the training set in order to construct and estimate *p*(*x*), we can then compute the log-likelihood of the kernel density estimate over the testing set:

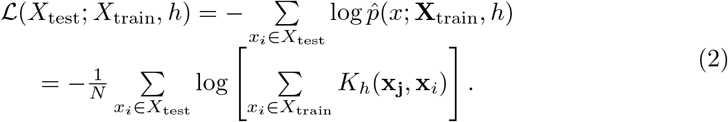

The bandwidth parameter *h* can therefore be determined by minimizing Eq. 2, which is a simple one-dimensional *f* function of *h*.

After all centers have been identified, labels are assigned to each data point by simply comparing the relative density gradients between a given point and the cluster centers. After labels have been assigned to each point, if, for a given neighborhood, 99 percent of the cells share the same cluster center, that cluster is considered “pure” and is added to the initial set of pure clusters.

Once HAL-x identifies the pure clusters, it performs a second, more rigorous validation of the point cluster assignment. We use two separate validation protocols to ensure that our initial neighborhoods have sufficiently high density and are not errant groupings that stem from a low density threshold. HAL-x uses the approximate nearest neighbors search algorithm a second time to define an extended neighborhood for each cluster center. This extended neighborhood includes points that are within a more relaxed density threshold than the one used for the approximate nearest neighbor search in the previous step. If there is a significant decrease in purity within the extended neighborhood, the conflicting clusters (representing the same density maxima) are merged to prevent unnecessary overlap.

### Supervised Linkage

After the initial “pure” clusters have been identified, HAL-x performs agglomerative clustering by linking clusters that are hard to distinguish. This supervised linkage between clusters is performed in the original high-dimensional feature space. This is done by training supervised learning classifiers (commonly random forests or support vector machines) to distinguish all *pairs of clusters* from each other. In other words, if there are *N* pure clusters, we train *N* (*N* − 1)/2 classifiers that distinguish between each pair of clusters. HAL-x then builds a minimally connected K-Nearest Neighbors (KNN) graph wherein each edge in the graph connected a cluster to the *K* clusters that are the most difficult to distinguish from itself (i.e. to the *K* clusters with the lowest out-of-sample accuracy) [26].

Using the KNN graph, HAL-x then performs a deeper sweep on each pair connected by an edge. This deeper sweep is performed with a larger ensemble of random forest classifiers built on bootstrapped samples of the data, thereby reducing the risk of overfitting. When HAL-x has completed this deeper sweep, the two clusters with the lowest edge-score (classification accuracy) are merged. The KNN graph and the labels are updated accordingly. This process (starting with a deep sweep of all clusters connected by an edge) is repeated until there is only a single cluster that contains all the data points (corresponding a perfect cv-score of 1).

The classification ensemble that generated the final merge is stored in the root node of a tree data structure. We chose a tree data structure so that a path can be traced between any pure cluster and the root. Excluding the pure clusters, which act as the leaves, each cluster is a parent for the two child clusters that were merged to create this parent cluster. The classifiers used to separate the child clusters are stored in the parent node, and each edge (separating a parent and a child node) is linked to the accuracy value (cv-score) that led to the merge.

### Prediction

Using the tree data structure created during the training process, HAL-x can ultimately predict the multiple cluster labels for millions of unreduced data points. For this purpose, the user selects a minimum score (cv-score), which corresponds to the accuracy of the cross-validated supervised models. Every predicted label will correspond to a cluster that can be separated from all other clusters with a mean accuracy score greater than the cv-score value. A higher cv–score will result in fewer clusters, whereas a lower cv–score will result in a large number of clusters that are only slightly differences from neighboring clusters (See Figure 3B-C). The default cv-score is 0.50: random classification. A cv-score of 0.50 will typically consider the full set of pure clusters when predicting the labels.

**Fig 3.**
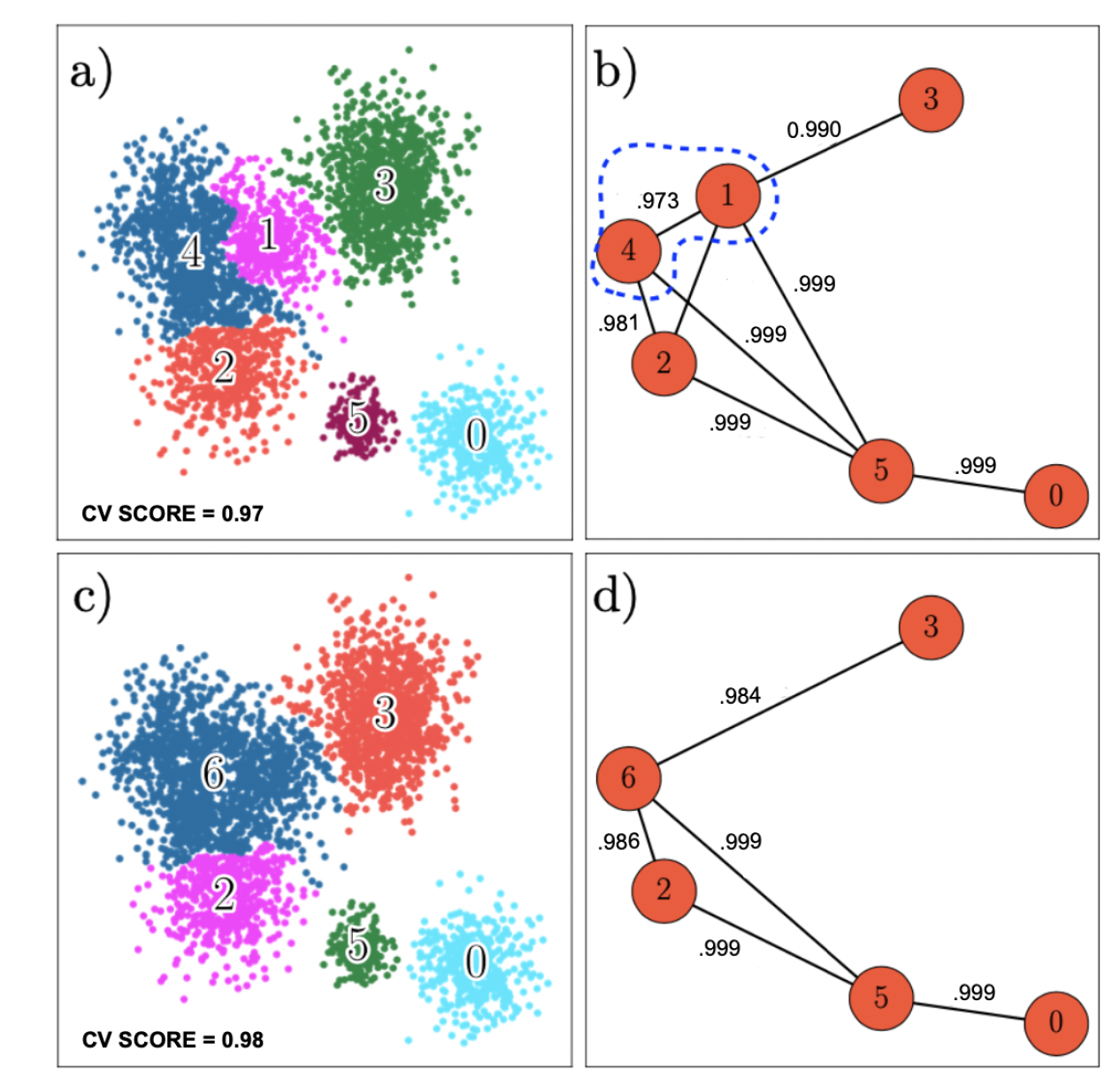
Clustering based on out-of-sample error. (a) Clusters generated with a cv-score of 0.97. (b) Graph illustrating out-of-sample error for each pair of clusters. There are no pairs of clusters with an out-of-sample error that is greater than 0.03 (accuracy value of 0.97). (c) Clusters generated with a cv-score of 0.98. Clusters 1 and 4 have been merged because the out-of-sample error was greater than (d) Graph illustrating accuracy for each pair of clusters. There are no pairs of clusters with an out-of-sample error that is greater than 0.02 (accuracy value of 0.98).

To make the predictions and thereby apply the clustering across all datasets, HAL-x first separates the data using the classifiers corresponding to the final merge that created the cluster containing all the data. Subsequent predictions are made using the classifiers from the children of the predicted cluster, further separating the data based on increasingly nuanced patterns. This process continues until the next predicted label would result in a cluster that is connected to the parent with an edge-score of less than cv-score. With the tree approach utilized here, a range of cv-score values can be efficiently checked for the same model, generating different sets of clusters depending on the level of population specificity desired by the user. Algorithm 1 highlights the simplicity of the HAL-x prediction technique via the cv-score parameter.

## Experimental Results

Here, we present the results of testing HAL-x on both synthetic and real-world datasets. First, we benchmarked HAL-x on the FlowCAP I dataset. We then generated multiple synthetic datasets containing gaussian populations. We used these datasets to compare the scalability of HAL-x to Parc, Phenograph, and HDBSCAN. We also tested HAL-x on a large, real-world mass cytometry dataset [4, 27], custom–generated at the NIH.

### Algorithm 1

HAL-x Prediction

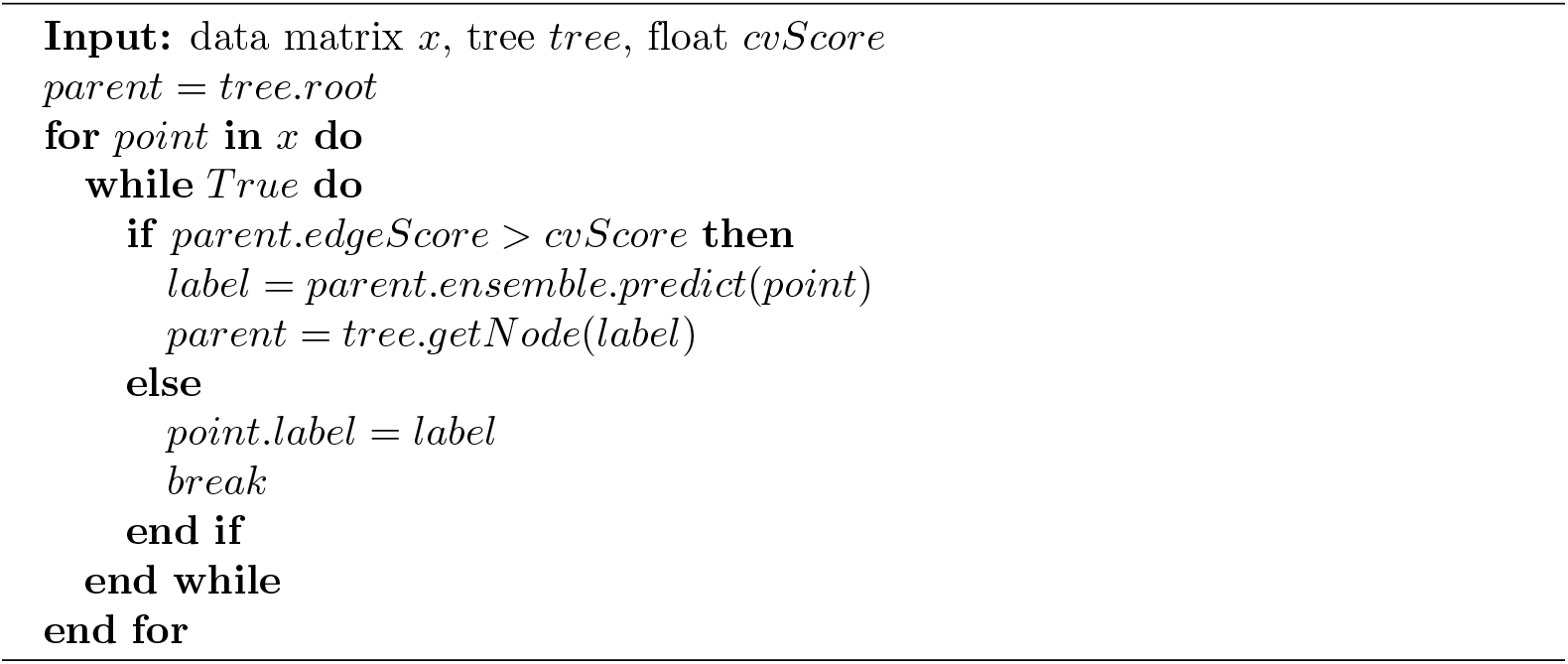

This dataset contains 25.9 million single-cells taken from healthy individuals and from patients with Lupus [28]: these samples were profiled with an antibody panel quantifying 34 surface epitopes. After clustering, we performed supervised classification to determine if HAL-x clusters could be used to discriminate clinical statuses from leukocyte profiles: this provides a foundation for future experimental work and drug/biomarker discovery.

### Levine Benchmark Datasets

We first tested HAL-x on two benchmark datasets from [5] to demonstrate the validity of HAL-x clusters: scalability is meaningless if the outputs are random. The first dataset is a flow cytometry dataset containing 81,747 human bone marrow mononuclear cells (BMMC) cells from 1 healthy tissue sample. These BMMC cells have 13 features corresponding to surface epitopes and a label corresponding to phenotype. The second dataset is a mass cytometry dataset containing 104,184 human bone marrow mononuclear cells (BMMC) cells from 2 healthy tissue samples. These BMMC cells have 32 features corresponding to surface epitopes and a label corresponding to phenotype. These phenotypes have been hand defined by flow cytometry experts and represent our general understanding of leukocytes in bone marrow. We used HAL-x to cluster these single–cell datasets and compared the multiclass F1-scores with those achieved by three other clustering algorithms: Parc, Phenograph, and HDBSCAN (Table 1).

**Table 1.**
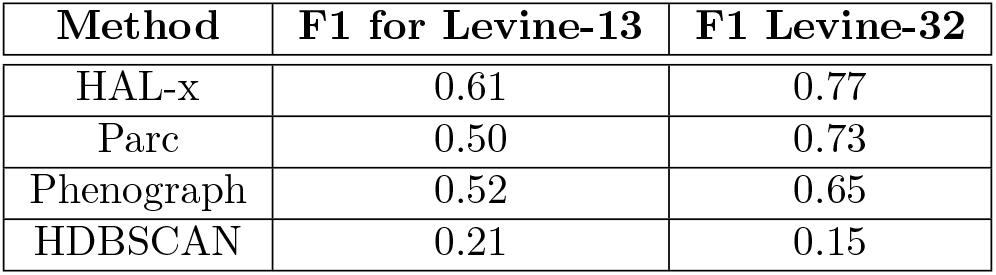
F1 Measure of clustering accuracy for diverse methods, applied onto CyTOF datasets from [5].

| Method | F1 for Levine-13 | F1 Levine-32 |
| --- | --- | --- |
| HAL-x | 0.61 | 0.77 |
| Parc | 0.50 | 0.73 |
| Phenograph | 0.52 | 0.65 |
| HDBSCAN | 0.21 | 0.15 |

We see from Table 1 that HAL-x significantly outperforms Parc, Phenograph, and HDBSCAN on both these datasets, showing HAL-X is competitive with existing techniques in terms of accuracy.

### Synthetic Datasets

HAL-x is designed as a convenient, accessible tool for biomedical researchers for this reason we wanted to assess how well HAL-x preformed on large datasets with limited computational resources. To do so, we generated a synthetic dataset with 10 millions data points, the typical size of a dataset in single-cell biology, and tested HAL-x on an 8-core laptop computer - the typical computing resources available to a bench researcher. High-performance computing was not used in this study. Within this computing environment, we compare HAL-x to the three algorithms widely used for fast clustering of high-dimensional data: Parc, Phenograph, and HDBSCAN.

We generated datasets with 10^3^, 10^4^, 10^5^, 10^6^, and 10^7^ synthetic data points, each with 30 features. The size and dimensions of these synthetic datasets were chosen to match the structure of mass cytometry data, which typically contains approximately 30 features (antibodies) and more than 10^7^ single cells. For HAL-x and HDBSCAN (algorithms which have prediction capabilities), we defined a 1 percent downsample to train on the datasets containing from 10^4^ to 10^6^ data points. For the dataset with 10^3^ points, we used a 10 percent downsample because 100 data points was insufficient for the perplexity of the fitSNE algorithm. We used a 0.5 percent downsample on the dataset with 10^7^ points so the memory required by the fitSNE algorithm would not exceed available resources in our computing environment. We do not perform any dimensionality reduction on the test dataset: without significant computing resources, this is typically infeasible for large datasets. Therefore, we are unable to reduce the training set for HDBSCAN, which requires both training and test data to have the same dimension. Parc and Phenograph cannot predict the cluster labels of new data points. For these methods, we attempted to learn the clustering model on the entire dataset (without downsampling). We used the default parameters for all the methods considered in this study.

In figure 4, we show the scalability of HAL-x compared to 3 algorithms that are commonly used for single-cell data clustering. We see that all 4 algorithms are relatively equal at 10^3^ and 10^5^. At 10^5^, Phenograph is significantly slower than HAL-x, HDBSCAN, and Parc. At 10^6^, both Parc and Phenograph exceed the available memory within our computing environment. HDBSCAN can train a model on 500,000 cells and predict on 10e7 cells, but the runtime is immensely slower than HAL-x. This observation is consistent with the predictive mechanism of HDBSCAN, which relies on nearest neighbour search rather than a set function learned by the supervised classifiers used in HAL-x.

**Fig 4.**
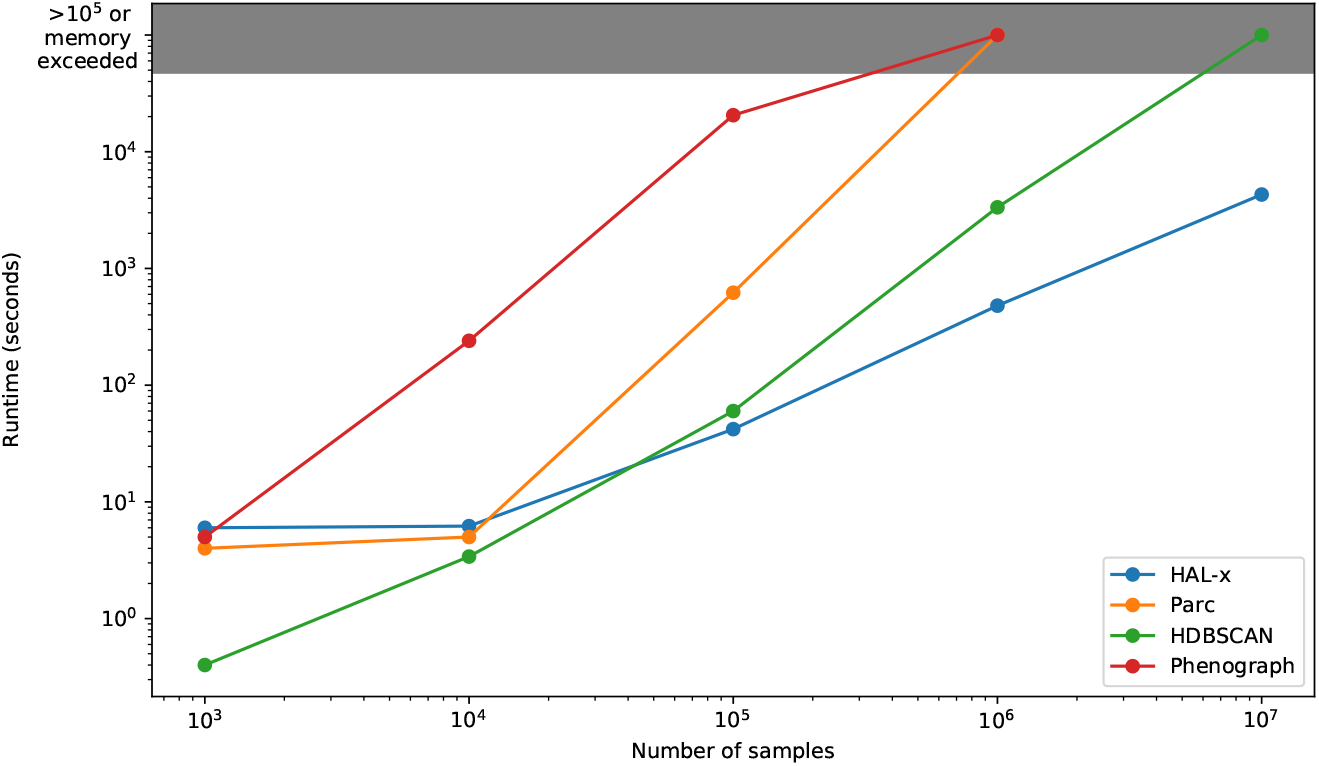
Comparison of run times for training and label prediction on synthetic datasets of increasing sizes. Graybox indicates when memory of laptop was exceeded on dataset.

### Lupus Dataset

Finally, to illustrate the scalability and tunability of HAL-x, we tested HAL-x on a real-world mass cytometry datasets containing 25.9 million individual cells analyzed from 75 peripheral blood mononuclear cells (PBMC) samples and 75 polymorphonuclear (PMN) samples. 49 of these samples were from Lupus patients and 26 of these samples were from healthy individuals. We designed a 38-marker mass cytometry panel with a focus on neutrophils as well as general phenotyping of blood leukocytes (see supplementary materials for experimental details).

At the onset of this study, we assumed no prior knowledge regarding which populations would constitute a good biomarkers for lupus and potential targets for immunotherapy. Thus, we sought to generate 5 separate sets of clusters, each with a different level of population specificity, ranging from very rare to quite broad.

In under 1 hour, HAL-x generated 5 separate clusterings for the 25.9 million cells. Figure 5B-C presents the linear relationship between the cv-score parameter and the number of clusters, demonstrating the ability of HAL-x to widely vary the number of clusters using the same model.

**Fig 5.**
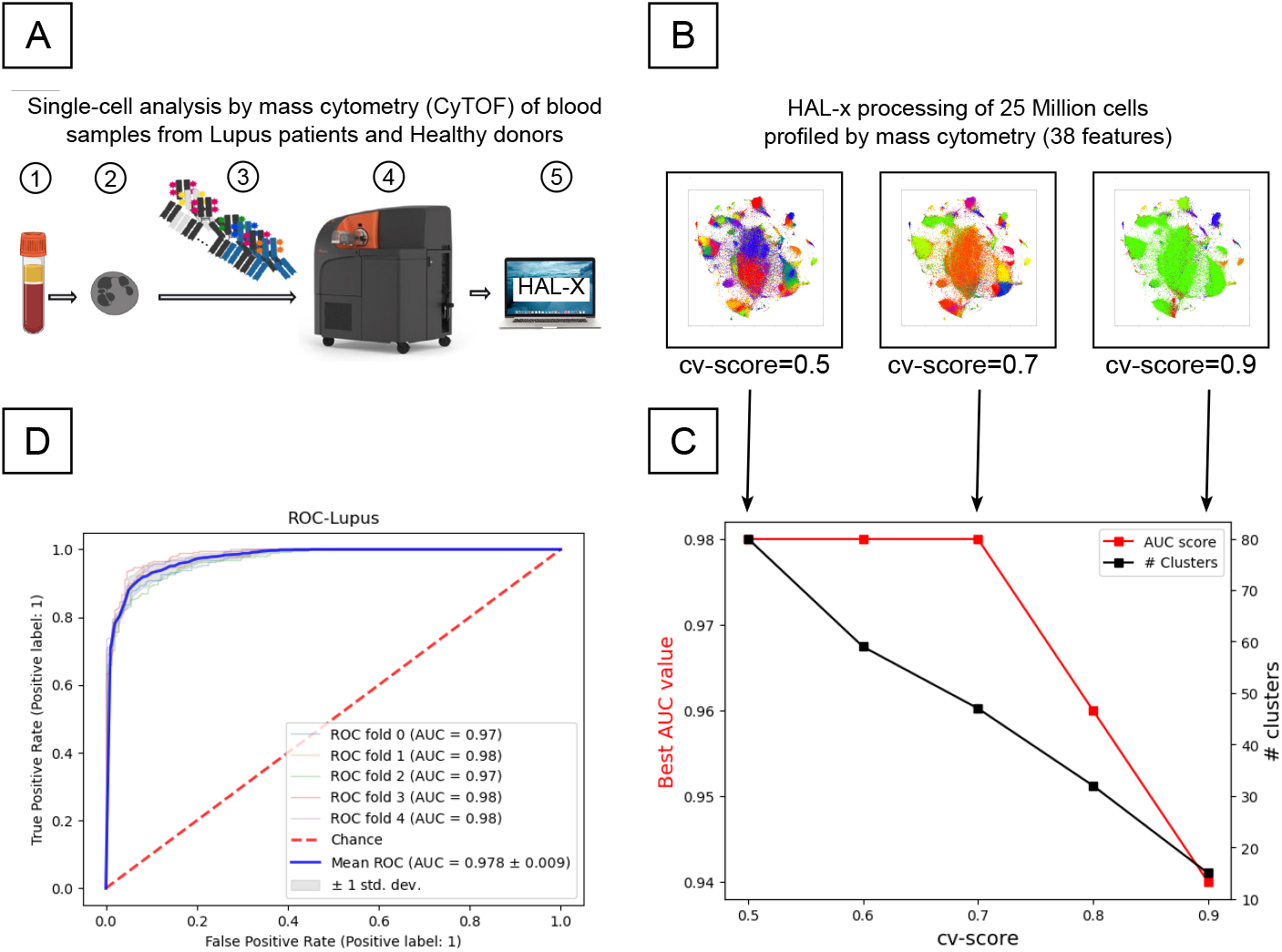
Analysis of Lupus patients’ blood profile using single-cell mass cytometry and HAL-x. (A) The pipeline for profiling immune cells from Lupus patients and healthy donors, beginning with blood collection, continuing with mass cytometry, and culminating with HAL-x processing. (B) t-SNE visualizations of clusterings generated by HAL-x which correspond to various levels of population specificity (cv-score). (C) A graph of AUC (*i.e.* Area under the Receiver Operating Curve (ROC)) vs. CV-score, showing that the most important clusters with discriminatory power are not observed for low CV-scores. (D) The ROC curve for the best cluster identified by HAL-x, separating individual neutrophils from Lupus and Healthy patients with an AUC of 0.82.

For each HAL-x cluster (across our 5 different clustering depths), we trained a random forest classifier to separate healthy cells and Lupus cells. We discovered multiple clusters of PBMC cells and neutrophils wherein the healthy cells and lupus cells were classified with an AUC score above 0.95. The altered phenotypes of these cell populations can be considered novel biomarkers for Lupus. These populations will be the subject of wet-lab experimentation seeking to highlight their functional relevance for Lupus pathology and to identify new molecular/cellular targets for lupus immunotherapies. Moreover, our results on the lupus dataset show the importance of varying the population specificity when searching for key populations in high-dimensional data. We observed here that several of the key cell populations were only identified when the cv-score was 0.7 or lower (deeper clustering). When the cv-score was higher (a more broad clustering), these important clusters are merged with other populations. This resulted in lower accuracy values and the loss of naunced phenotypes which may have clinical signficance. Our results show the importance of testing multiple clustering with varying depths. For example, if we had fixed the cv-score at 0.8, we would have missed several biomarkers and lost valuable information about the cellular landscape. HAL-x offered both the robustness, tunability and scalability to match the biological complexity of our Lupus dataset.

### Conclusion

The emergence of inexpensive platforms for large-scale single-cell analysis in biomedical applications is presenting new challenges to researchers attempting to interpret these immense, high-dimensional datasets. HAL-x integrates density clustering (reduced data) and supervised classification (unreduced data) to generate clusters that can be projected across tens of millions of single-cells.

The success of HAL-x on our lupus datasets shows the importance of clustering algorithms that: 1) can easily predict points for new data and 2) can interact with raw data. These are vital components in the time/memory efficiency achieved by HAL-x. Moreover, the simple HAL-x prediction algorithm (Algorithm 1) allows for one model to generate multiple sets of labels corresponding to varying levels of population specificity - a valuable advantage for biomedical researchers who need to cluster as deep as possible to identify key populations (biomarkers) of biological relevance and as shallow as possible to retain statistical significance in these biomarkers.

The success of HAL-x clustering when used for supervised classification tasks (*i.e.* identifying a novel cluster of neutrophils that distinguishes lupus patients from their healthy counterparts) suggests that our approach has a promising role in biomarker derivation and drug discovery. For example, the trends within specific subpopulations can be used to predict targets for new Lupus immunotherapies. More generally, the ability of HAL-x to generate a clustering model that can be propagated and tuned across multiple datasets will standardize the ability of biological researchers to analyze vast amount of data and to communicate their results.

## Acknowledgements

PM and AD were supported by Simons Investigator in the Mathematical Modelling of Living Systems Grant, and NIH Grant No. 1R35GM119461. This research was supported in part by the Intramural Research Program of the NIH (MJK and GA-B research groups). This project was initiated with a seed grant to GA-B and PM from the Gordon and Betty Moore foundation and Research Corporation through the Scialog program.

## Supplementary Methods

### CyTOF mass cytometric profiling of Lupus blood samples

#### Acquisition of single–cell measurements

Fresh Blood samples from 50 Lupus patients and 25 healthy volunteers were collected. Samples were immediately processed to separate low-density and high-density neutrophils (LDN and NDN) using Histopaque gradient. LDN and NDN fractions were then collected and washed in prewarmed complete RPMI immediately prior to cell-surface staining and acquisition. All reagents were from Fluidigm unless otherwise noted. Cells were first incubated with a 5-*μ*M solution of cisplatin in PBS to mark dead cells. The cells were then washed and resuspended in Maxpar Cell Staining Buffer (MCSB) (Fluidigm). Human Fc receptor blocking solution (BioLegend) was added to each sample and incubated for 10 minutes at room temperature. All metal-labeled antibodies were purchased from Fluidigm. Biotin anti–human CD115 (BioLegend) was detected with Qdot-streptavidin conjugate (Thermo Fisher Scientific). A table of the panel of antibodies used for our mass cytometric measurements can be found in S1. Antibodies were diluted in MCSB at concentrations validated for minimal channel spillover, added to each sample, and incubated for 30 minutes at room temperature. Cells were washed twice with MCSB and then fixed for 15 minutes in a 1.6% solution of paraformaldehyde (MilliporeSigma) in PBS. Cell intercalation solution was prepared by adding Cell-ID Intercalator-Ir into Maxpar Fix and Perm Buffer (both from Fluidigm) to a final concentration of 125 nM. Cells were incubated with cell intercalation solution overnight at 4°C and then washed once with MCSB and Maxpar Water twice. Cells were finally resuspended in a water solution containing EQ Four Element Calibration Beads (Fluidigm) immediately prior to CyTOF data acquisition at a cell concentration of 10^6^cells/ml (¡300 events per second) and filtered into cell trainer cap tubes. Data were acquired on a Helios Mass Cytometer (Fluidigm) as previously described [27].

#### Data preprocessing

Raw mass cytometric data were normalized with the Normalizer algorithm as recommended by the software developers. Data were first gated as live singlet cells using (EQ4–cisplatin) and (CD45^+^DNA(2n)) manual gates (Supplemental Figure S2) and exported as Python dataframes. These dataframes were then processed with a custom–programmed pipeline in Python, based on an implementation of HAL-X.

### Complexity for generating multiple clusterings

The scalability of HAL-x is facilitated by the supervised classifiers used to merge the clusters. We simply use the trained models to predict the cluster labels on the raw data. For multiple sets of clusters, we use classifiers at different levels of the hierarchy. The worst-case time complexity for prediction via a trained random forest is *O(υ×t)* where *υ* is the number of variables (features) and *t* is the number of trees in the forest. The worst-case time complexity for prediction via a trained SVM is *O(υ×fs)* where *υ* is the number of variables and *s* is the number of support vectors. With this sub-exponential runtime, HAL-x can easily predict labels for datasets with over 25 million cells, as we show in this report.

#### Algorithm 2

HAL-x

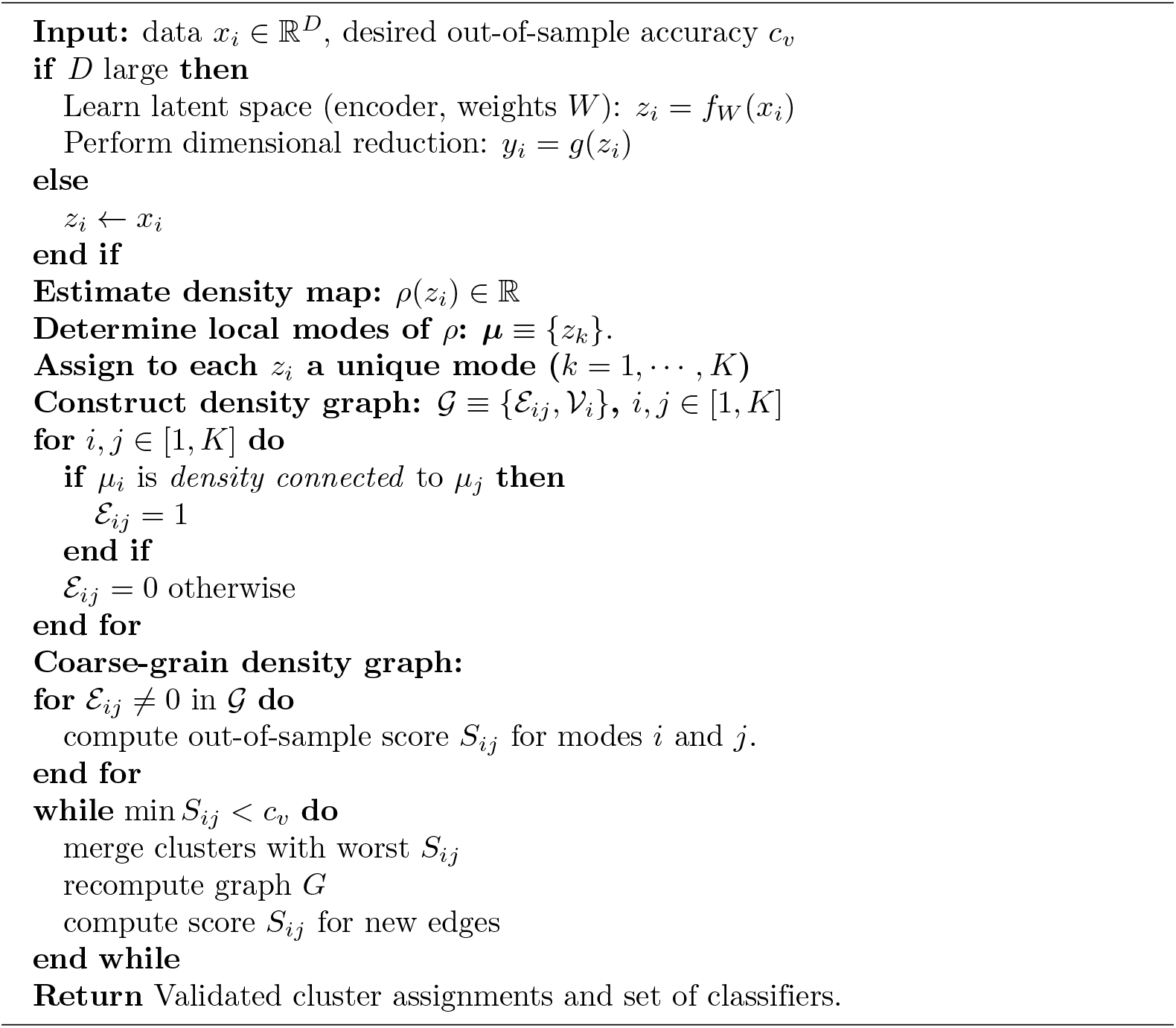

### HAL-x algorithm

In Algorithm 2, we present the pseudocode representation of HAL-x model generation (end-to-end).

### Supervised classification of single cells

First, we used random sampling to balance the data contained in each cluster, ensuring an equal amount of cells from Lupus patients and from healthy individuals. For each cluster, we used a random forest classifier (with bootstrapping) to predict the clinical status of the individual cells (Lupus or healthy).

---

**Fig S1.**
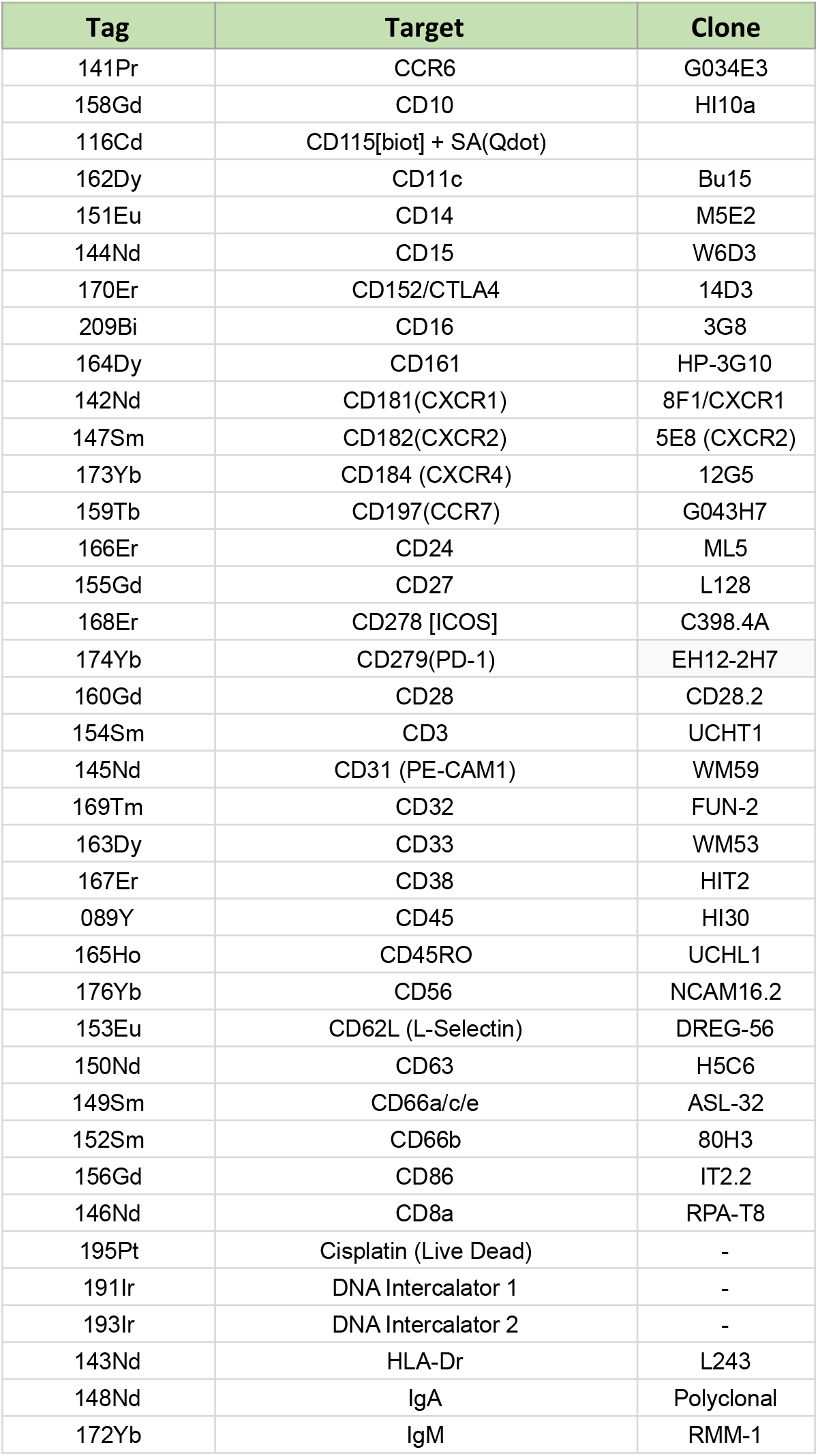
Panel of metal-coupled antibodies used to profile the blood samples from Lupus patients and healthy donors.

**Fig S2.**
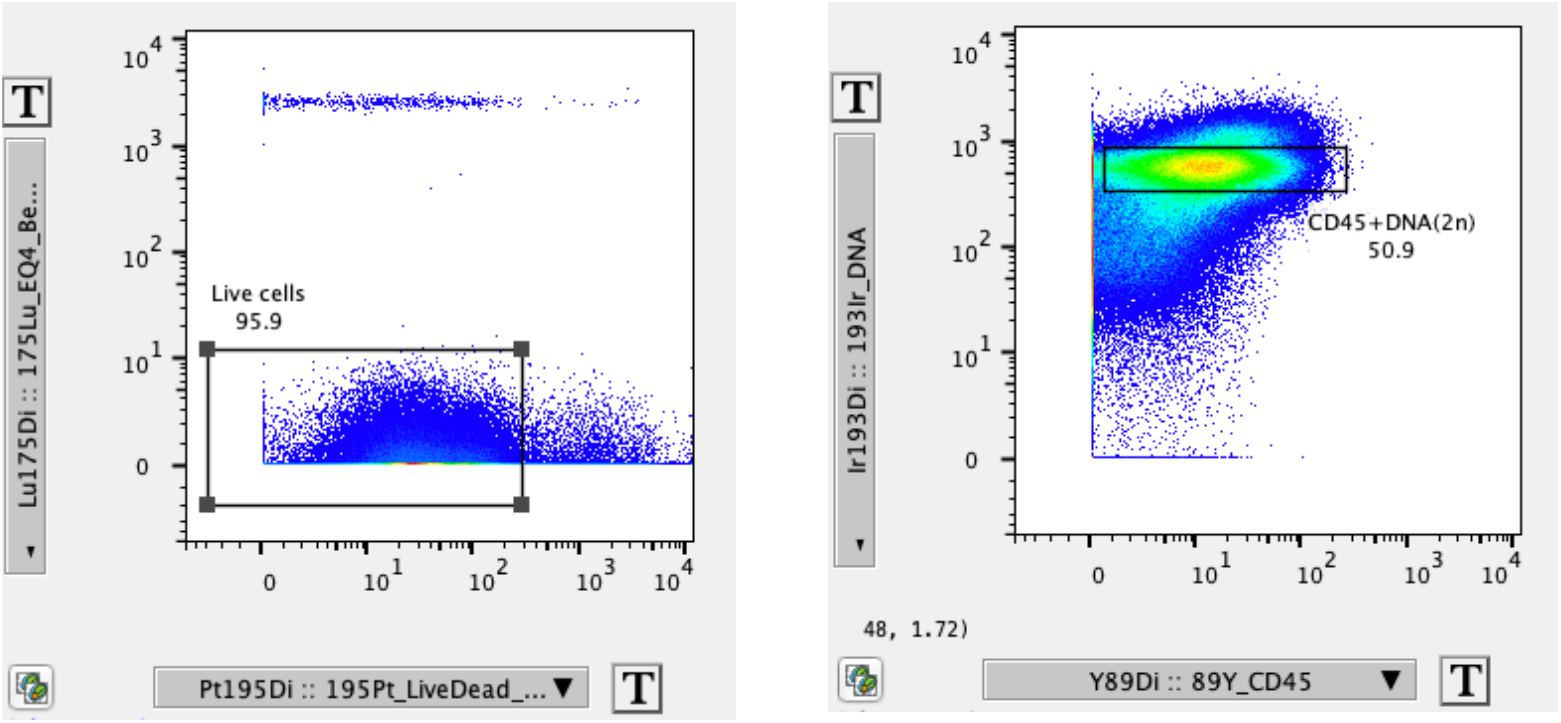
Raw .fcs file processing and gating strategy. Left: gating for live cells (excluding Cisplatin-positive dead cells and EQ4 calibration beads. Right: gating for CD45^+^DNA(2n) singlet leukocytes.

